# Mesenchymal stromal cell chondrogenesis under ALK1/2/3-specific BMP inhibition: A revision of the prohypertrophic signalling network concept

**DOI:** 10.1101/2024.03.12.584629

**Authors:** Solvig Diederichs, Simon I. Dreher, Sarah-Anna Nüesch, Sven Schmidt, Christian Merle, Wiltrud Richter

## Abstract

**Background:** In vitro chondrogenesis of mesenchymal stromal cells (MSCs) driven by the essential chondro-inducer transforming growth factor (TGF)-β is instable and yields undesired hypertrophic cartilage predisposed to bone formation in vivo. TGF-β can non-canonically activate bone morphogenetic protein-associated ALK1/2/3 receptors. These have been accused of driving hypertrophic MSC misdifferentiation, but data remained conflicting. We here tested the antihypertrophic capacity of two highly specific ALK1/2/3 inhibitors – compound A (CompA) and LDN 212854 (LDN21) – in order to reveal potential prohypertrophic contributions of these BMP/non-canonical TGF-β receptors during MSC in vitro chondrogenesis.

**Methods:** Standard chondrogenic pellet cultures of human bone marrow-derived MSCs were treated with TGF-β and CompA (500 nM) or LDN21 (500 nM). Daily 6-hour pulses of parathyroid hormone-related peptide (PTHrP[1–34], 2.5 nM, from day 7) served as potent antihypertrophic control treatment. Day 28 samples were subcutaneously implanted into immunodeficient mice.

**Results:** All groups underwent strong chondrogenesis, but GAG/DNA deposition and *ACAN* expression were slightly but significantly reduced by ALK inhibition compared to solvent controls along with a mild decrease of the hypertrophy markers *IHH-, SPP1-*mRNA, and Alkaline phosphatase (ALP) activity. When corrected for the degree of chondrogenesis (*COL2A1* expression), only pulsed PTHrP but not ALK1/2/3 inhibition qualified as antihypertrophic treatment. In vivo, all subcutaneous cartilaginous implants mineralized within 8 weeks, but PTHrP pretreated samples formed less bone and attracted significantly less haematopoietic marrow than ALK1/2/3 inhibitor groups.

**Conclusions:** Overall, our data show that BMP-ALK1/2/3 inhibition cannot program mesenchymal stromal cells toward stable chondrogenesis. BMP-ALK1/2/3 signalling is no driver of hypertrophic MSC misdifferentiation and BMP receptor induction is not an adverse prohypertrophic side effect of TGF-β that leads to endochondral MSC misdifferentiation. Instead, the prohypertrophic network comprises misregulated PTHrP/hedgehog signalling and WNT activity, and a potential contribution of TGF-β-ALK4/5-mediated SMAD1/5/9 signalling should be further investigated to decide about its postulated prohypertrophic activity. This will help to successfully engineer cartilage replacement tissues from MSCs in vitro and translate these into clinical cartilage regenerative therapies.

## Background

Articular cartilage tissue lacks the intrinsic capacity to regenerate and thus, traumatic cartilage injuries pose a high risk to develop into osteoarthritis which is not curable. Cell and tissue engineering therapies based on autologous chondrocytes have been developed and optimized to treat focal cartilage injuries but disadvantages are obvious. Cartilage biopsy for chondrocyte harvest is highly invasive and the limited tissue resources make in vitro chondrocyte expansion and a second surgery for cell implantation necessary. As a promising alternative, mesenchymal stromal cells (MSCs) are widely studied, since they are capable to differentiate into chondrocytes, can be isolated in large amounts from regenerating tissues like bone marrow and allow a one-step surgery. While MSC in vitro differentiation into chondrocytes has long been achieved [1, 2], the intrinsic propensity for endochondral misdifferentiation into hypertrophic chondrocytes was not fully overcome. Expressing Indian hedgehog (IHH), type X collagen, alkaline phosphatase (ALP), osteopontin (OPN or SPP1) and other typical markers, hypertrophic chondrocytes exhibit an undesirable mineralization activity and a strong tendency to degenerate and induce bone formation at ectopic sites in vivo [2–4]. In our recent break-through study, a novel heparin-equipped biomaterial augmented with transforming growth factor (TGF)-β allowed stable chondral lineage commitment of MSCs in an in vivo setting [5]. In vitro, however, our continuing inability to induce stable chondral MSC differentiation shows that our understanding of the specific molecular events that are necessary to stabilize the chondrocyte phenotype still remains incomplete.

Two important regulators of endochondral MSC misdifferentiation are the WNT signalling network and the feedback signalling loop between parathyroid hormone-related peptide (PTHrP) and hedgehog. Pulsed PTHrP application and WNT inhibition foster MSC chondrogenesis and can decrease but not fully suppress hypertrophy to the low levels observed in articular chondrocytes [6–10]. Thus, further prohypertrophic signalling appears to be involved [6].

The bone morphogenetic protein (BMP)-associated SMAD signal transducers, SMAD1, SMAD5, and SMAD9 have often been accused of prohypertrophic activity in mice and humans, but their role during human MSC in vitro chondrogenesis remains surprisingly contradictive. SMAD1/5/9 proteins become activated during TGF-β-driven MSC chondrogenesis, and cell-autonomous BMP expression and signalling is one putative source [11, 12]. BMPs activate SMAD1/5/9 via recruitment of the activin receptor-like kinases ALK1, ALK2, ALK3 and ALK6 [13, 14]. In turn, the ALK4 and ALK5 receptors are activated by canonical TGF-β signalling and phosphorylate primarily prochondrogenic and antihypertrophic SMAD2 and SMAD3 proteins (Fig. 1). Importantly, TGF-β can further cross-activate SMAD1/5/9 in many cell types by various mechanisms [11, 15–17]. These include i) TGF-β cross-binding to ‘BMP’-ALK receptors [15, 18, 19], ii) cross-phosphorylation of SMAD1/5/9 by ALK4/5 [15, 20], and iii) formation of mixed receptor-regulated SMAD complexes [21, 22]. Surprisingly, it remains still unclear whether the potentially prohypertrophic SMAD1/5/9 activation during MSC in vitro chondrogenesis is part of a BMP-induced or TGF-β-derived action via ALK1/2/3 receptors or may occur as inevitable downstream effect of ALK4/5.

**Fig. 1.**
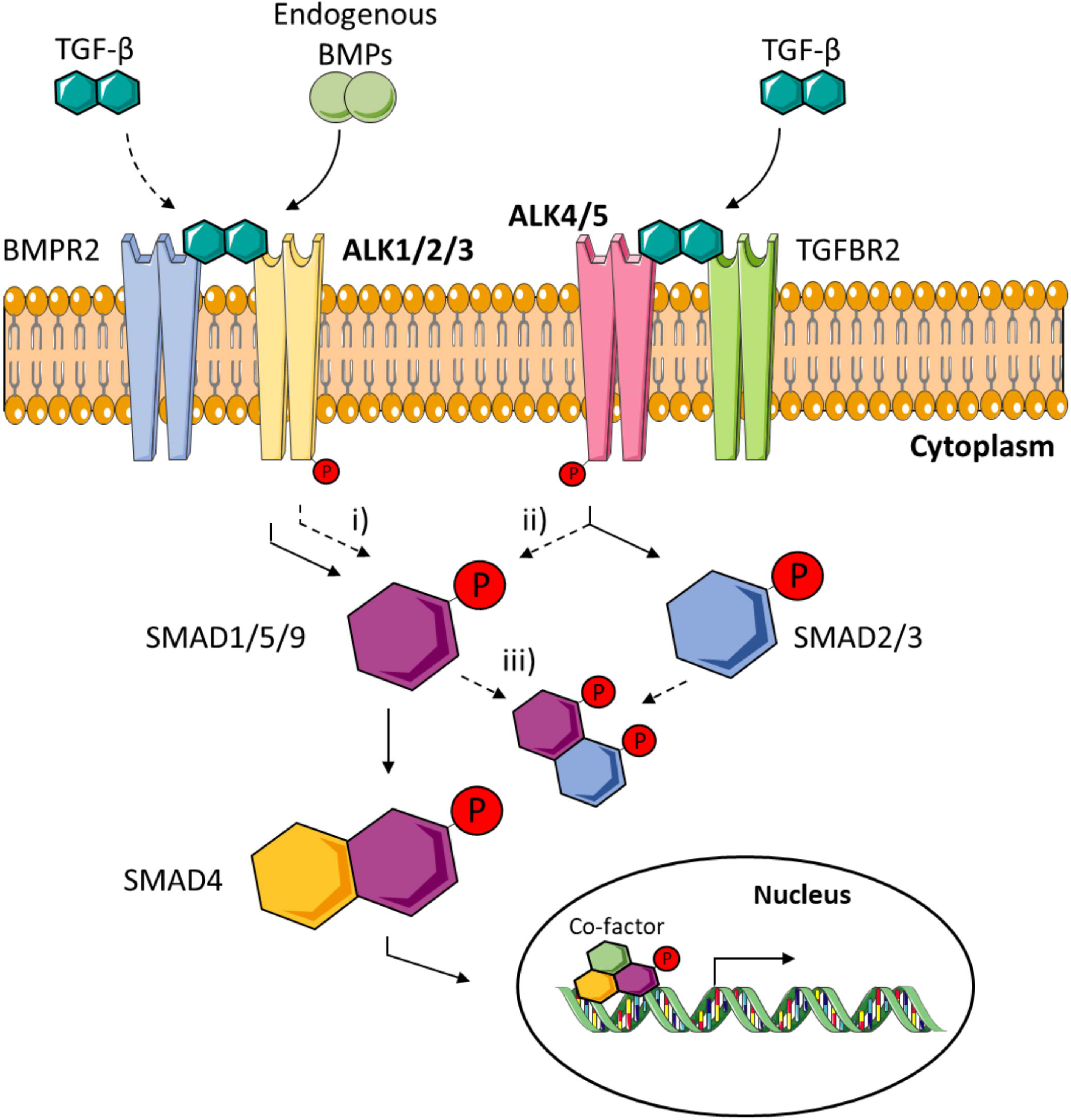
TGF-J3-mediated cross-activation of SMAD1/5/9. Cross-activation of SMAD1/5/9 can occur by i) TGF-B cross-binding to ’BMP’-ALK receptors [15, 18, 19], ii) by phosphorylation of SMAD1/5/9 by ALK4/5 [15, 20], or iii) by formation of mixed receptor-regulated SMAD complexes [21, 22]. This figure was generated using Servier Medical Art, provided by Servier, licensed under a Creative Commons Attribution 3.0 unported license (https://smart.servier.com).

In several studies ‘BMP’-ALK1/2/3 inhibitors were applied with the intend to suppress SMAD1/5/9 activation and modify the outcome of human MSC in vitro chondrogenesis. An antihypertrophic activity of the small-molecule inhibitor dorsomorphin was suggested by Hellingman et al., but we observed opposingly that weaker hypertrophic markers in dorsomorphin-treated MSCs were largely a consequence of less chondrogenesis [11, 12]. While in these studies, the high dorsomorphin dose may have allowed off-target SMAD2/3 inhibition, more recent studies dosed novel inhibitors with a strong BMP receptor bias in a range effective for ALK1/2/3 but not for ALK4/5; but results were again conflicting. Occhetta et al. reported that ALK1/2/3 inhibition with a proprietary inhibitor from Novartis called compound A sufficiently reduced hypertrophy markers in differentiating MSCs to install a stable chondrocyte phenotype and thereby attributed a prohypertrophic role to SMAD1/5/9 activation by ALK1/2/3 [24]. In contrast, Franco et al. reported that the ALK1/2/3 inhibitor LDN 193189 did not reduce hypertrophy unless applied in a high ALK4/5-affecting dose (1000 nM LDN19) which also reduced proteoglycan deposition, thus indicating a negligible role of ALK1/2/3 for hypertrophic degeneration of MSC-derived chondrocytes. This calls for a direct comparative evaluation of such novel small-molecule inhibitors in comparison to a potent antihypertrophic positive control to decipher the role of ALK1/2/3-mediated SMAD1/5/9 activation during MSC in vitro chondrogenesis.

The aim of this study was therefore to assess whether activation of ALK1/2/3 receptors by endogenous BMPs or the essential chondro-inducer TGF-β exerts prochondrogenic and/or adverse prohypertrophic side effects leading to endochondral MSC misdifferentiation via downstream activation of SMAD1/5/9. We blocked ALK1/2/3 during MSC chondrogenesis with the two strongest selective inhibitors available, compound A and LDN 212854 [23, 24], and used pulsed PTHrP application as antihypertrophic control treatment. Chondrogenic differentiation was determined by quantitation of proteoglycan and type II collagen deposition, and chondrocyte hypertrophy by specific marker expression as well as in vivo mineralization and bone formation in an ectopic implantation model.

## Methods

### BMP-activity assay

The BMP-activity assay was carried out as described before [25]. In brief, C2C12 cells were cultured in DMEM high-glucose, 10% fetal calf serum (FCS), 1% penicillin/streptomycin. After 24 hours, cells were stimulated with 150 ng/mL BMP2 for 3 days in the presence of 500 nM compound A (CompA), LDN 212854 (LDN21) or the solvent DMSO at equal concentration. Cells were washed with PBS and lysed with 1% Triton-X-100 at 4°C. Cell lysates (100 µL) were incubated with 100 µL of substrate solution (1 mg/mL p-nitrophenyl phosphate in 0.1 M glycine (Carl Roth, Germany), 1 mM MgCl_2_, and 1 mM ZnCl_2_ (all from Sigma Aldrich, Germany), pH 9.6). Absorbance was recorded at 405/490 nm (Sunrise, Tecan, Switzerland), enzyme activity was referred to a p-nitrophenol-derived standard curve (Sigma Aldrich, Germany) and normalized to protein levels determined in the same lysate by Pierce BCA Protein Assay Kit (Thermo Scientific, USA).

### Isolation and expansion of MSCs

Fresh bone marrow aspirates from patients, age 35-82 (mean 55) years undergoing total hip replacement surgery were acquired with informed written consent of the patients. Samples were obtained after approval by the Ethics Committee on Human Experimentation of the Medical Faculty of Heidelberg University and in agreement with the Helsinki Declaration of 1975 in its latest version. MSCs were isolated, as described [2]. Briefly, samples were subjected to Ficoll Paque Plus density gradient centrifugation. The mononuclear cell fraction was seeded in expansion medium consisting of high-glucose Dulbecco’s modified Eagle’s medium (DMEM), supplemented with 12.5% FCS, 2 mM L-glutamine, 1% nonessential amino acids, 0.1% β-mercaptoethanol (all from Gibco, Life Technologies, Germany), 1% penicillin/streptomycin (Biochrom, Germany), and 4 ng/mL of recombinant basic fibroblast growth factor (Miltenyi Biotec, Germany). Cells were cultured for 2 passages at a seeding density of 5×10 cells/cm.

### Chondrogenic differentiation

MSCs were cultured as 3D pellets (2.5×10 or 5×10 cells/pellet) in chondrogenic medium (DMEM high-glucose (Gibco, Life Technologies, Germany), 0.1 μM dexamethasone, 0.17 mM ascorbic acid 2-phosphate, 5 μg/mL transferrin, 5 ng/mL selenous acid, 1 mM sodium pyruvate, 0.35 mM proline (all from Sigma Aldrich, Germany), 1% ITS+ premix (Corning, Germany) 1% penicillin/streptomycin, 10 ng/mL recombinant human TGF-β1 (Pepro-Tech or Miltenyi, Germany) for up to 4 weeks at 37°C and 6% CO_2_. Cultures with 5×10 cells/pellet received daily medium exchange with or without 6 hours of prior PTHrP(1-34) treatment (2.5 nM in water, Bachem, Germany) from day 7 onward as described before [9] (Fig. 2). Chondrogenic medium of cultures with 2.5×10 cells/pellet was supplemented with the BMP inhibitor compound A (compA, 500 nM in 0.02% DMSO, days 0-28, Novartis, Switzerland) as described before [24], LDN 212854 (LDN21, 500 nM in 0.02% DMSO, day 0-28, Sigma Aldrich, Germany), SB-431542 (SB43, 10 μM, Merck Millipore, Darmstadt, Germany) or the respective solvent. Medium was exchanged 3 times a week (Fig. 2).

**Fig. 2.**
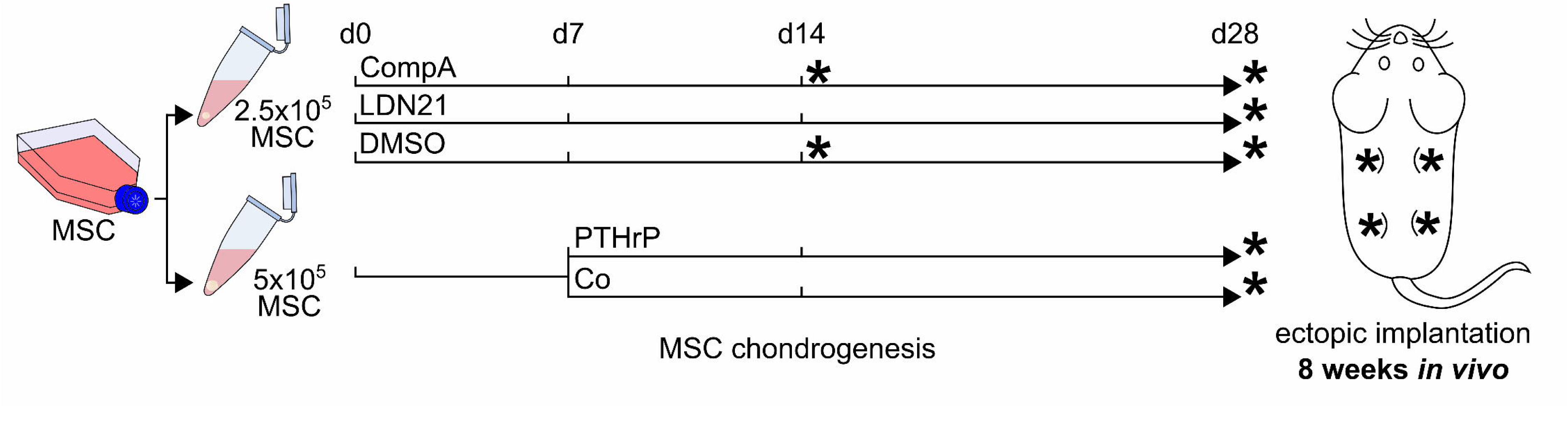
Study design: In vitro treatment and time of ectopic implantation in vivo. MSCs in passage 2 were subjected to chondrogenic induction in the presence of the BMP inhibitors CompA (500 nM), LDN21 (500 nM) or with DMSO used as control. Medium was exchanged three times a week. The PTHrP pulse group was treated with PTHrP (2.5 nM) for the last 6h before daily medium exchange from day 7 onward. Its control group (Co) also received daily medium exchanges starting on day 7.

### Quantitative real-time RT-PCR

Total RNA was extracted from three pooled pellets per donor population, group, and time point by guanidinium isothiocyanate/phenol extraction (peqGOLD Trifast, Peqlab, Germany), which was then reverse transcribed into cDNA using the reverse transcriptase Omniscript (Qiagen, Germany) and oligo(dT) primers. The expression levels of individual genes of interest were determined by quantitative real time PCR (qPCR, Roche Diagnostics, Germany or Stratagen, USA), primer pairs used for amplification are listed in Table 1. Genes were rated as expressed when gel electrophoresis of PCR amplificates showed a clear band at the correct height. Gene expression levels were calculated using the ΔCt method where the arithmetic mean expression of the reference genes *CPSF6* and *HNRPH1* was subtracted from the Ct value of the gene of interest. Percent reference gene was calculated as percentage of 1.8^(-ΔCt) and referred to solvent-treated (DMSO for CompA and LDN21 groups) or daily medium exchanged groups (Co for the PTHrP group) to give the fold-change value.

**Table 1.**
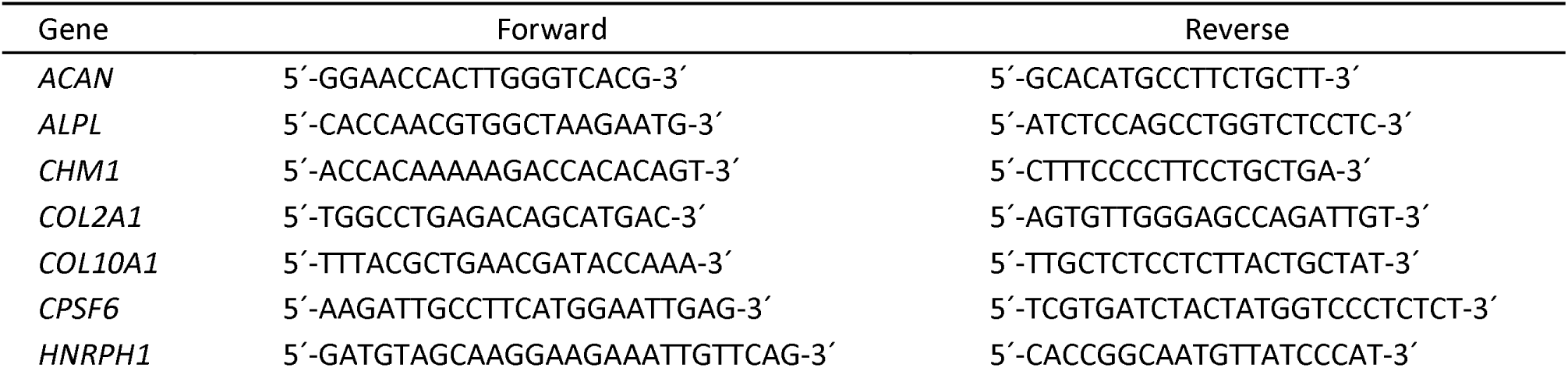
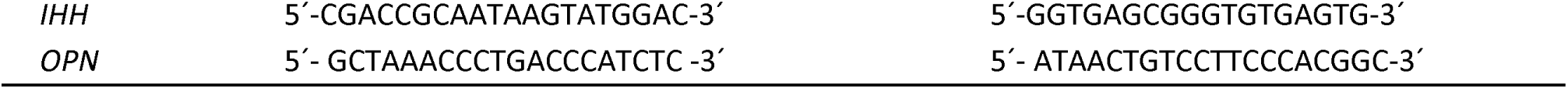
Primer pairs for qRT-PCR analysis in alphabetical order.

### ALP activity

Culture supernatants conditioned for 2 days were collected, pooled, and 100 µL conditioned media were incubated with 100 µL of p-nitrophenyl phosphate substrate solution. ALP activity was determined as described above.

### Western Blotting

Collagen extraction was performed as previously described [11]. In brief, one pellet per donor population and group was digested with pepsin solution containing 2.5 mg pepsin/mL in 0.5 M acetic acid and 0.2 M NaCl for 16 hours. The pH was neutralized with 1 M Tris base prior to extraction of the collagens with 4.5 M NaCl (overnight at 4°C, both Roth, Karlsruhe, Germany). After centrifugation, the pellets were resuspended in 400 μL precipitation buffer (0.1 M Tris base, 0.4 M NaCl) and the collagens precipitated for 4 hours at -20°C with 100% ethanol. After centrifugation, the pellets were resuspended in lysis buffer (50 mM Tris, 150 mM NaCl, 1% Triton X-100).

For SMAD protein detection, whole cell lysates were prepared from two pellets per group using Phosphosafe Extraction Reagent (Merck Millipore, Darmstadt, Germany) containing 1 mM Pefablock SC (Sigma Aldrich, St. Louis, MO, USA). After centrifugation, supernatants were mixed with Laemmli buffer (33.2 % (w/v) glycerol (Carl Roth, Karlsruhe, Germany), 249 mM Tris-HCl pH 6.8, 8.0 % (w/v) sodium-dodecyl sulfate (SDS), 0.02 % bromophenol blue (all Sigma Aldrich, St. Louis, MO, USA) and boiled.

Cell or collagen lysates were separated by denaturing SDS-PAGE (6% gels for collagens, 10% for SMADs) and proteins blotted onto a nitrocellulose membrane (Amersham™, GE Healthcare, Chalfont St Giles, United Kingdom). Membranes were cut horizontally to detect proteins of different sizes. For collagen detection, the lower part of the membrane was incubated with mouse anti-human type X collagen antibody (X53, 1:500, Quartett, Germany) and the upper part with mouse anti-human type II collagen antibody (II-4C11, 1:1000, ICN Biomedicals, USA). For SMAD detection, upper membrane parts were first probed with rat monoclonal anti-pSMAD1/9 antibody (pS463/pS465, pS465/pS467, 1:1,000, #562508, BD Biosciences, East Rutherford, NJ, USA), or rabbit monoclonal anti-pSMAD2 (S465/467, 1:500, clone 138D4). For total SMAD detection, membranes were re-stained with rabbit monoclonal anti-SMAD1/5 (SMAD1: 1:500, ab33902; SMAD5: 1:1,000, ab40771; both Abcam, Berlin, Germany) or rabbit monoclonal anti-SMAD2/3 (1:250, clone D7G7, both Cell Signaling Technologies, Danvers, MA, USA). The lower membrane parts were probed with mouse monoclonal anti-β-actin (1:10,000, clone AC-15, GTX26276, GeneTex, Irvine, CA, USA). Proteins were visualized with HRP-conjugated goat anti-rat antibody (1:1,000, HAF005, Bio-Techne, R & D Systems, Minneapolis, MN, USA), peroxidase-coupled goat anti-mouse antibody (1:10,000, #115-035-071), or peroxidase-coupled goat anti-rabbit antibody (1:10,000, #111-035-046, both Jackson ImmunoResearch Laboratories, West Grove, PA, USA), using the ECL detection system (Roche, Germany).

### In vivo mineralization model

All animal experiments were approved by the local animal experimentation committee and carried out in accordance with European Laboratory Animal Science guidelines. The mineralization and bone formation activity of the pellets was tested in the standard subcutaneous model in immunocompromised mice. After 28 days of chondrogenic culture, two pellets per donor population (n=4) and group were implanted into paravertebral subcutaneous pouches (one in a cranial pouch, one in a caudal pouch) of 10-12 week old female SCID mice (CB17/Icr-*Prkdc* /Crl, Charles River, Sulzfeld, Germany) to account for the biological variability and allow significant conclusions. For CompA and the respective control group, also day 14 pellets were tested. Four subcutaneous pouches were prepared per mouse and one pellet per pouch was implanted. In total, 46 pellets that were pretreated as indicated were implanted into 12 mice. Two additional control pellets generated under standard chondrogenic conditions (no DMSO, thrice per weak medium exchange) that were not included in the analyses were implanted in one mouse. None of the mice exhibited any adverse reactions to the presence of the implants and had to be excluded from the study according to the defined exclusion criteria defined in the animal license (e.g., behavioural abnormalities, infections, fever, severely disturbed general condition). Explants were harvested after 8 weeks and stored at -80°C.

### Microcomputed tomography

Micro-CT evaluation was used as primary outcome to assess in vivo mineralization. Explants were scanned with a Sky-Scan 1076 in vivo X-ray microcomputed tomograph against air in a custom-made humid chamber to prevent drying during scanning. Settings were: no filter, 49 kV, 250 µA, voxel size 9 µm, exposure time 900 ms, frame averaging 3. Pictures were recorded every 0.42 degrees rotation step through 360 degrees. NRecon® software (version 1.6.3.2, Skyscan) was used for reconstructing 3D pictures. CTAn® software was used for calculating the total volume (TV) and the mineralized volume (MV) in mm of scanned tissues. Background was defined by unspecific signals in scans against air, and an 8-bit grey level threshold of 25 was set to discriminate signals from background. To define a threshold value for tissue mineralization, comparative scans of mineralized explants and osteochondral biopsies were performed in PBS vs. air. Non-mineralized tissue was defined by being indiscernible from aqueous surroundings, and a grey level above 66 in scans against air was determined as threshold for mineralized tissue. This threshold was validated in scans of osteochondral biopsies. Mineralized tissue volume (MV) of pellets was referred to the total volume (TV) to calculate mineralization levels (MV/TV).

### Histology

Pellets and explants were fixed for 2 hours in 4% formaldehyde. Explants (but not cultured samples) were partially decalcified for 2h with Bouin’s solution. Samples were dehydrated in a graded isopropanol series and paraffin-embedded. 5 µm tissue sections were deparaffinized, rehydrated and stained with 0.2% (w/v) Safranin O (Fluka, Sigma Aldrich, Germany) in 1% acetic acid, or alizarin red-S (0.5% in water) using Certistain Fast Green (Merck, Germany, 0.04% (w/v) in 0.2% acetic acid) as counterstain, following standard protocols. An overview staining was performed by haematoxylin and eosin staining according to standard protocols. Immunohistology was performed as described [2]. Briefly, rehydrated 5 µm tissue sections were treated with 4 mg/mL hyaluronidase in PBS, pH 5.5 and 1 mg/mL pronase (both from Roche Diagnostics, Germany), blocked with 5% bovine serum albumin (Sigma Aldrich, Germany) and stained with anti-human type II or I collagen antibody (1:1,000, clone II-4C11, ICN Biomedicals, Germany; or Abcam, Germany), followed by biotinylated goat anti-mouse antibody (1:500, Dianova, Germany) and streptavidin alkaline phosphatase fast red (Roche Diagnostics).

### Histological scoring

Haematoxylin-eosin stained paraffin sections of explants were scored by six blinded scorers to identify and semi-quantify bone formation as well as haematopoiesis. Haematopoiesis was defined as areas with marrow-cavity-like structures with the presence of haematopoietic cells and lymphocytes and evaluated in relation to the whole section. Bone was defined as dense, plain structure with osteocytes and evaluated in relation to the whole section. The semi-quantitative score ranged from 0 to 3 with the following definition: (0) non-existent, (1) low: 0-30% of the section area, (2) intermediate: 30-60%, (3) high: 60-100% of the scored section area. Mean scores for each treatment regimen were calculated for each scorer for statistical evaluation.

### Statistics

Results are shown as median values and are depicted as boxplots, with each box representing the interquartile range (IQR) extending between the 25th and 75th percentiles and lines inside the boxes representing the median. Whiskers extend to minimum and maximum values, outliers (between 1.5xIQR and 3xIQR) are depicted as 1, and extreme values (>3xIQR) as ◊. Statistical significance between two groups was calculated using Mann-Whitney U test. Where indicated, the *p* value was adjusted for multiple comparisons using Bonferroni correction. For time courses, data are given as the mean ± standard deviation. A probability value of *p*≤0.05 was considered statistically significant. All statistical tests were calculated with SPSS 25.0 (IBM, Germany).

## Results

### Slight antichondrogenic effects of ALK1/2/3-biased BMP-receptor inhibitors

Before the ALK1/2/3 inhibitors CompA and LDN21 were applied during MSC chondrogenesis, their dose-dependent action was tested in a biofunctional BMP activity assay in C2C12 cells. Both inhibitors significantly suppressed the BMP2-induced activation of alkaline phosphatase (ALP) in C2C12 cells at concentrations above 50 nM and lost their effect below (Suppl. Fig S1). The commonly chosen concentration of 500 nM was applied for both inhibitors in further experiments [23, 24].

The design of the main study, assessing prochondrogenic activity and antihypertrophic effects of ALK1/2/3-biased BMP inhibition versus daily pulsed PTHrP treatment on cartilage neogenesis of MSCs, is detailed in Fig. 2. For both study arms, cell count of pellets, compound concentration, timing of treatment and in vivo implantation were strictly adjusted to the parameters reported in previous studies to obtain the most promising results for each study arm and for reproducibility reasons [9, 24]. After 4 weeks of in vitro chondrogenesis, a strong proteoglycan and type II collagen-rich matrix had accumulated under all conditions according to toluidine blue (Fig. 3A), safranin O (Fig. 3B) and type II collagen staining (Fig. 3C) of histological sections. As expected, pellets in the PTHrP study arm groups appeared larger than pellets in the ALK inhibition study arm, which is consistent with higher starting cell numbers in these pellets and a more frequent medium exchange. A trend to less safranin O staining under ALK1/2/3 inhibition with CompA and LDN21 contrasted the more intense proteoglycan staining of PTHrP pulsed pellets compared to the corresponding controls (Fig. 3A,B). In line, quantitation of the GAG/DNA content compared to the respective control groups reflected significantly lower values under ALK1/2/3-selective inhibition while PTHrP pulsed pellets contained significantly more GAG/DNA (median 1.5-fold) compared to DMSO or medium controls (Fig. 3D). Along, aggrecan (*ACAN*) mRNA levels were significantly reduced in CompA or LDN21 groups and higher expressed by trend under PTHrP pulse treatment. Expression of chondromodulin 1 (*CHM1*) was also significantly reduced by CompA while it was significantly elevated after pulsed PTHrP treatment (Fig. 3E). Expression of *COL2A1* was generally less affected except a significant reduction in the CompA group (Fig. 3E). Type I collagen deposition remained unaffected by any treatment (suppl Fig. S2). Overall, although permitting strong cartilage neogenesis from MSCs in vitro, ALK1/2/3 inhibition showed some antichondrogenic effects while PTHrP pulse treatment was prochondrogenic. Thus, ALK1/2/3 signalling contributed to prochondrogenic TGF-β signalling, albeit only to some degree.

**Fig. 3.**
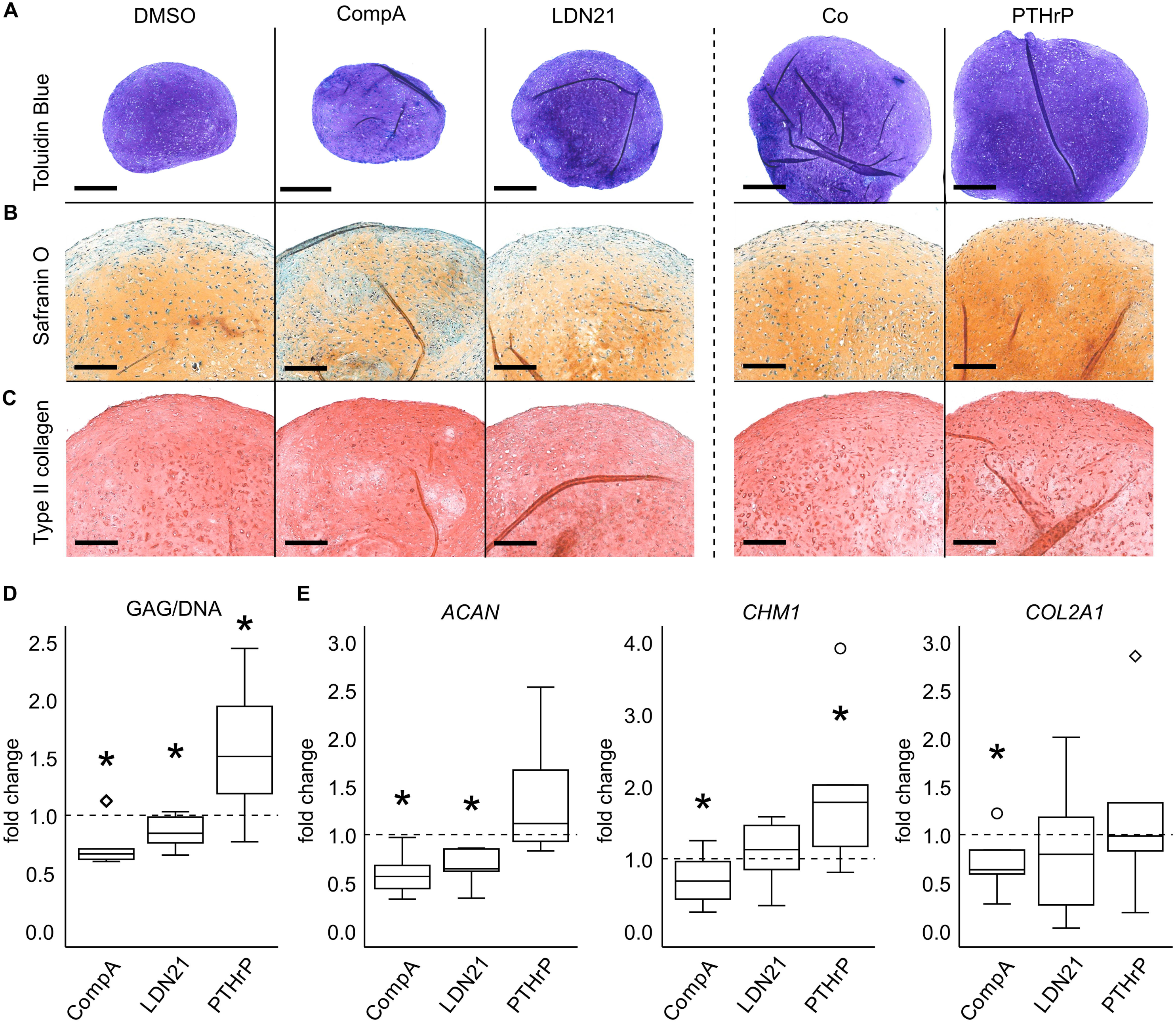
Effect of BMP pathway inhibition or PTHrP pulse treatment on cartilage matrix deposition and expression of differentiation markers. MSCs were subjected to chondrogenic induction for 28 days in the presence of CompA, LDN21 or received PTHrP pulse treatment. Paraffin sections were stained with (A) toluidine blue and (B) safranin O/fast green to visualize proteoglycan deposition or for (C) type II collagen by immunohistochemistry. Shown is one representative experiment of n=6 MSC donors. Scale bars represent 500 µm for toluidine blue overviews and 200 µm for magnified pictures. (D) Quantification of GAG and DNA content (n=S MSC donors). (E) Expression levels of indicated genes with HNRPH1 and CPSF6 as reference (n=6 donors). In D­ E, respective control cultures were set to 1 (dashed lines). Boxes represent the interquartile range (IQR, 25th to 75th percentiles), lines inside the boxes represent the median, whiskers extend to minimum and maximum values. Outliers are depicted as o and extreme values as◊. * p:::;;0.05 Mann-Whitney U test compared to respective controls.

### Reduction of endochondral markers by ALK1/2/3 inhibition and PTHrP pulse treatment

Next focus was the impact of ALK1/2/3 inhibition on the lineage choice of MSCs. A common set of hypertrophic (*COL10A1* and *IHH*) and osteogenic (*SPP1*=*OPN* and *ALPL*) marker genes was investigated on day 28 of chondrogenesis (Fig. 4). In contrast to Occhetta et al., only weak effects on lineage choice were seen for CompA treatment, which did not significantly lower *COL10A1, IHH,* and *OPN* expression, while *ALPL* expression was significantly but only slightly reduced (Fig. 4A). LDN21 treatment significantly reduced *COL10A1*, *IHH* and *ALPL* expression while *OPN* was only affected by trend (Fig. 4A). However, when referred to *COL2A1* expression to correct for reduced chondrogenesis, none of the regulations remained significant (suppl. Fig. S3), demonstrating that weaker endochondral signals under BMP inhibition were largely accessory to less chondrogenesis. In contrast, PTHrP pulse treatment induced much stronger suppression of *IHH*, *OPN* and *ALPL* and resulted in the lowest median values for *COL10A1* expression of all 3 groups (Fig. 4A). These effects were robust against referring to *COL2A1* expression (suppl. Fig. S3).

**Fig. 4.**
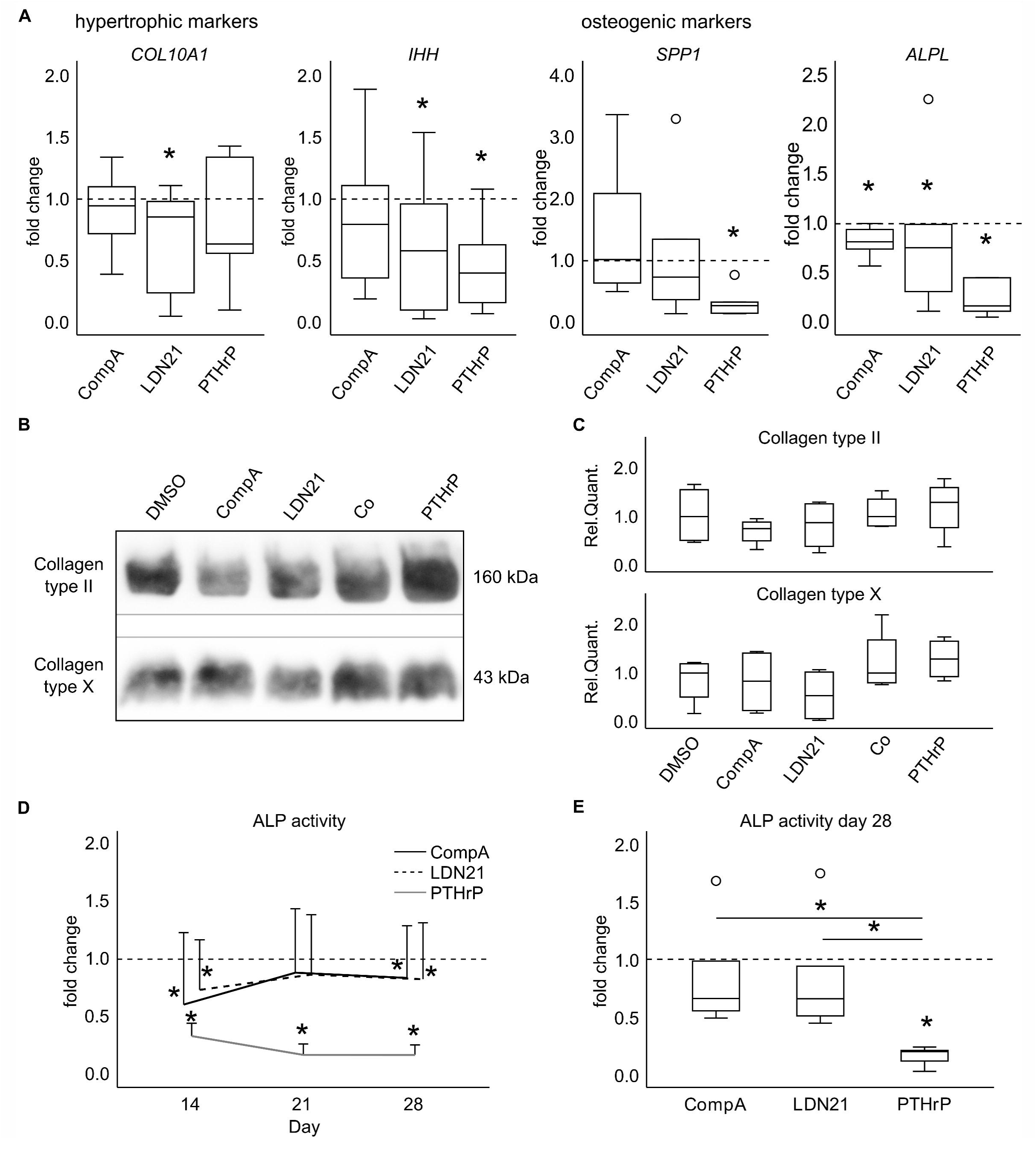
Effect of BMP pathway inhibition or PTHrP pulse treatment on endochondral marker expression. MSCs were subjected to chondrogenic induction for 28 days in the presence of CompA, LDN21 or received PTHrP pulse treatment. Expression levels of (A) hypertrophic and osteogenic marker genes, referred to the respective control cultures (dashed lines, n=6 donors). (B) Western blotting of type II collagen and type X collagens extracted from 1 pellet per group. Representative results from one of four experiments (n=4 donors) are shown. Please note, that pepsin digestion used for collagen extraction degrades all other proteins, so that typical loading controls cannot be provided. The image was cropped for conciseness as indicated and full-length blots are presented in supplementary Fig. S7. (C) Densitometric quantification of the Western blots in (B) (n=4). (D) Alkaline phosphatase (ALP) activity was determined in culture supernatants in weekly intervals (n=6 donors). Data are referred to respective control cultures (dashed lines) and shown as mean and standard deviation. (E) ALP activity on day 28 referred to respective control cultures (dashed lines). Box plots were generated as described in Fig. 3. * p:50.05 Mann-Whitney U test with Bonferroni correction compared to respective controls and between treatments.

In line with mRNA data, little effect was obvious on type X and type II collagen deposition by ALK1/2/3 inhibition according to densitometry of Western blots (Fig. 4B,C). Also pulsed PTHrP treatment was not antihypertrophic enough to considerably affect type X collagen deposition in line with our previous data [10].

Although ALP enzyme activity in the culture supernatants was significantly reduced by ALK1/2/3 inhibition when compared to controls (Fig. 4D, dashed line), effects were relatively low. In contrast, strong suppression of ALP activity was observed under pulsed PTHrP treatment. At termination of culture, pulsed PTHrP treatment resulted in a significantly stronger reduction of mineralization-relevant ALP activity than ALK1/2/3 inhibition with either compound (Fig. 4E). For ALK1/2/3 inhibition, ALP activity effects were lost when referred to *COL2A1* expression, but remained for pulsed PTHrP treatment after correction (suppl. Fig. S3). A full silencing like in articular chondrocytes was not obtained as observed before [6]. Taken together, PTHrP pulse treatment but not ALK1/2/3 inhibition induced broad antihypertrophic effects, documenting the responsivity of the MSC donor populations. However, under ALK1/2/3 inhibition, the reduced chondrocyte hypertrophy appeared only as a consequence of reduced chondrogenesis, suggesting that ALK1/2/3 signalling is rather irrelevant for hypertrophy during TGF-β-driven MSC chondrogenesis, and hypertrophy appears as inevitable downstream effect of ALK4/5 and/or other pathways.

### Cartilage mineralization after ectopic implantation

Next, we addressed whether the partial antihypertrophic effects of ALK1/2/3 inhibition or PTHrP pulse application influenced cell fate permanently or whether phenotype stabilization was only transient and depended on continuous inhibitor supply. Day 28 cartilage samples were implanted into subdermal pouches on the back of immunodeficient mice for 8 weeks. Histology showed that the proteoglycan content of explants was strongly reduced compared to pre-implantation samples according to toluidine blue and safranin O staining, with the best retention observed in PTHrP pulsed pellets (Fig. 5A). Obviously, cells were insufficiently primed regarding self-sustaining growth factor production to maintain a high level of cartilage matrix production after discontinuation of TGF-β treatment. Alizarin red staining revealed strong mineralization in all groups including pulsed PTHrP treatment where less bone and more calcified cartilage seemed to occur (Fig. 5A). For quantitation of total explant volume (TV) and mineralized tissue volume (MV), samples were scanned against air by microcomputed tomography (micro-CT). 3D reconstruction of data and a heat map of mineralization of central tissue sections confirmed strong tissue mineralization in explants from all groups (Fig. 5B, Suppl. Fig. S4). Tissue of treatment groups retained a similar TV than respective controls in vivo (Fig. 5C) and no differences in MV/TV were obvious between treatment groups and respective controls (Fig. 5D). Thus, neither in vitro pretreatment was able to reduce in vivo tissue mineralization demonstrating that a desired irreversible cell fate decision and induction of a self-sustaining permanent articular chondrocyte phenotype was not achieved by any of them. While this may have been expected for BMP-ALK inhibition, pulsed application of the hedgehog inhibitor PTHrP is our currently most effective antihypertrophic treatment that is capable of some decoupling of chondrogenesis from hypertrophy markers. However, like with WNT inhibition in vitro, a permanent fate shift similar to that of heparin-coupling of TGF-β in vivo was not possible and the reduction of hypertrophy remained dependent on inhibitor supply.

**Fig. 5.**
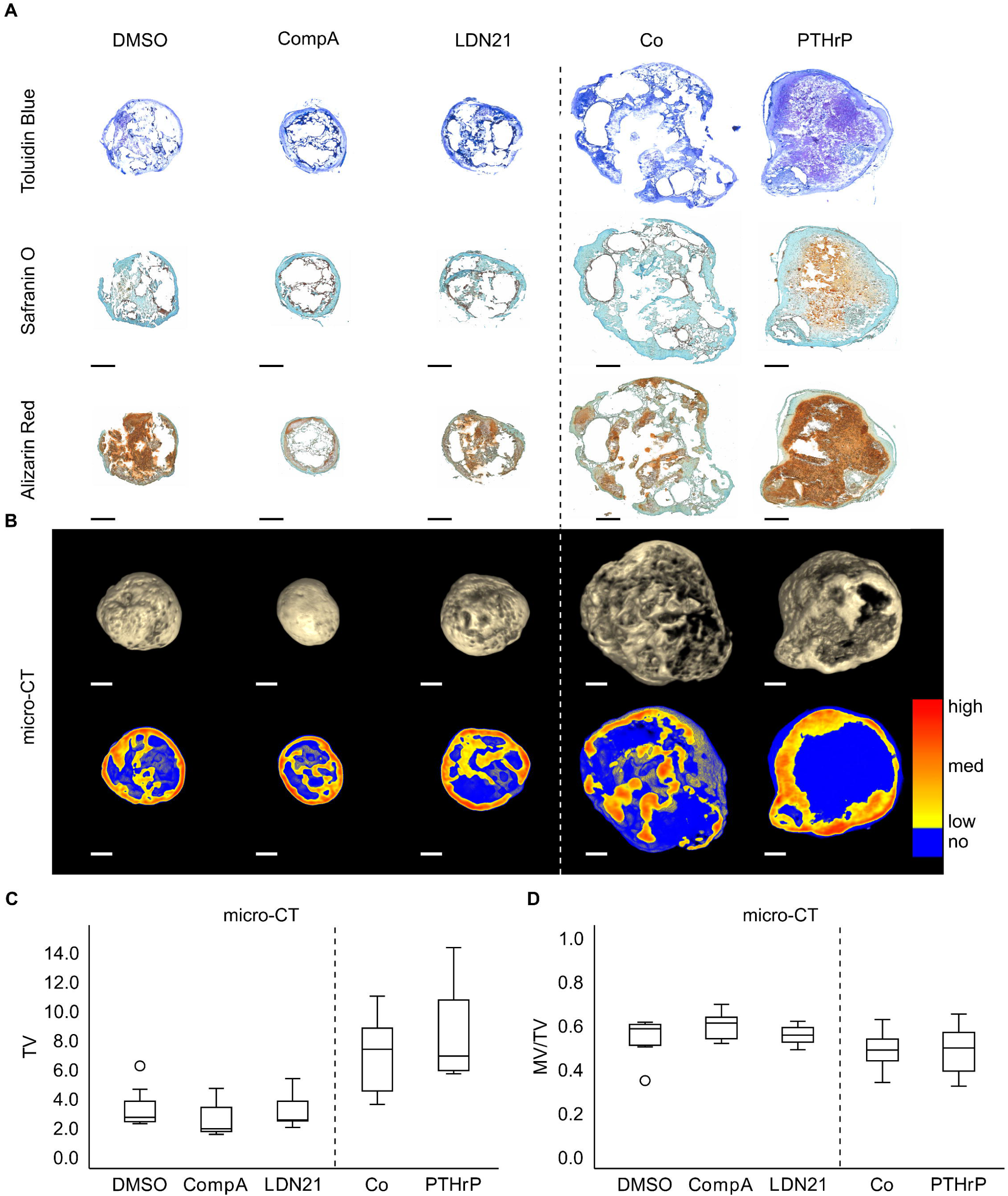
Effect of BMP pathway inhibition or PTHrP pulse treatment on mineralization of MSC-derived cartilage after 8 weeks in vivo. MSCs were subjected to chondrogenic induction in the presence of CompA, LDN21 or received PTHrP pulse treatment. After 28 days, samples were implanted in subcutaneous pouches on the back of SCIO mice for 8 weeks. Paraffin sections of explants were stained with (A) toluidine blue, safranin 0/fast green to visualize proteoglycan deposition or alizarin red to visualize mineralization. Shown is one representative experiment based on n=?-8 implants from n=4 MSC donors. (B) Mineralization analysis by micro-CT. Full 3D reconstructions (top row) and central sections {bottom row) with a colour-gradient (yellow­ low to red-high) representing mineralization levels. Non-mineralized tissue was visualized in blue.. All samples are shown in suppl. Fig. S4. Scale bars represent 500 µm. (C) Mineralized volume (MV) in mm3 referred to total volume {TV) in mm3 (n=?-8 experiments from n=4 MSC donors). Box plots were generated as described in Fig. 3. Mann-Whitney U test compared to respective controls.

### Bone formation and attraction of haematopoietic marrow in vivo

A still remaining question was whether the explants showed a similar degree of bone formation and attraction of haematopoietic marrow depending on in vitro pretreatment. Thus, HE-stained histological sections were scored for the presence of bone and haematopoietic tissue by several blinded observers (Fig. 6). Significantly less bone and haematopoietic tissue was obtained in the PTHrP pulse group compared to its control indicating some enduring effect of this treatment after implantation (Fig. 6B). In contrast to Occhetta et al., ALK1/2/3 inhibitor pretreatment with CompA had no obvious effects on in vivo neocartilage calcification and bone formation of parallel samples and the same was observed for LDN21 treatment (Fig. 6B). Since Occhetta et al. implanted precultured MSC pellets treated with CompA after 2 weeks of chondrogenic preculture in vitro, we added experiments with 4 additional MSC donor populations with their study design but obtained the same results as after 4 weeks of preculture: strong loss of proteoglycans from cartilage, strong mineralization and bone formation with no significant differences from DMSO controls (Suppl. Fig. S5, S6). In conclusion, ALK1/2/3 blocking during MSC chondrogenesis is not able to shift MSCs into the chondral lineage to produce a permanent neocartilage tissue resistant to in vivo degeneration, mineralization and bone formation. Thus, neither BMP nor TGF-β-associated ALK1/2/3 activation was relevant for endochondral MSC differentiation, suggesting that either ALK4/5-mediated cross-activation of SMAD1/5/9 (Fig 1, i), mixed SMAD complexes (Fig1, ii) and/or other pathways are more relevant.

**Fig. 6.**
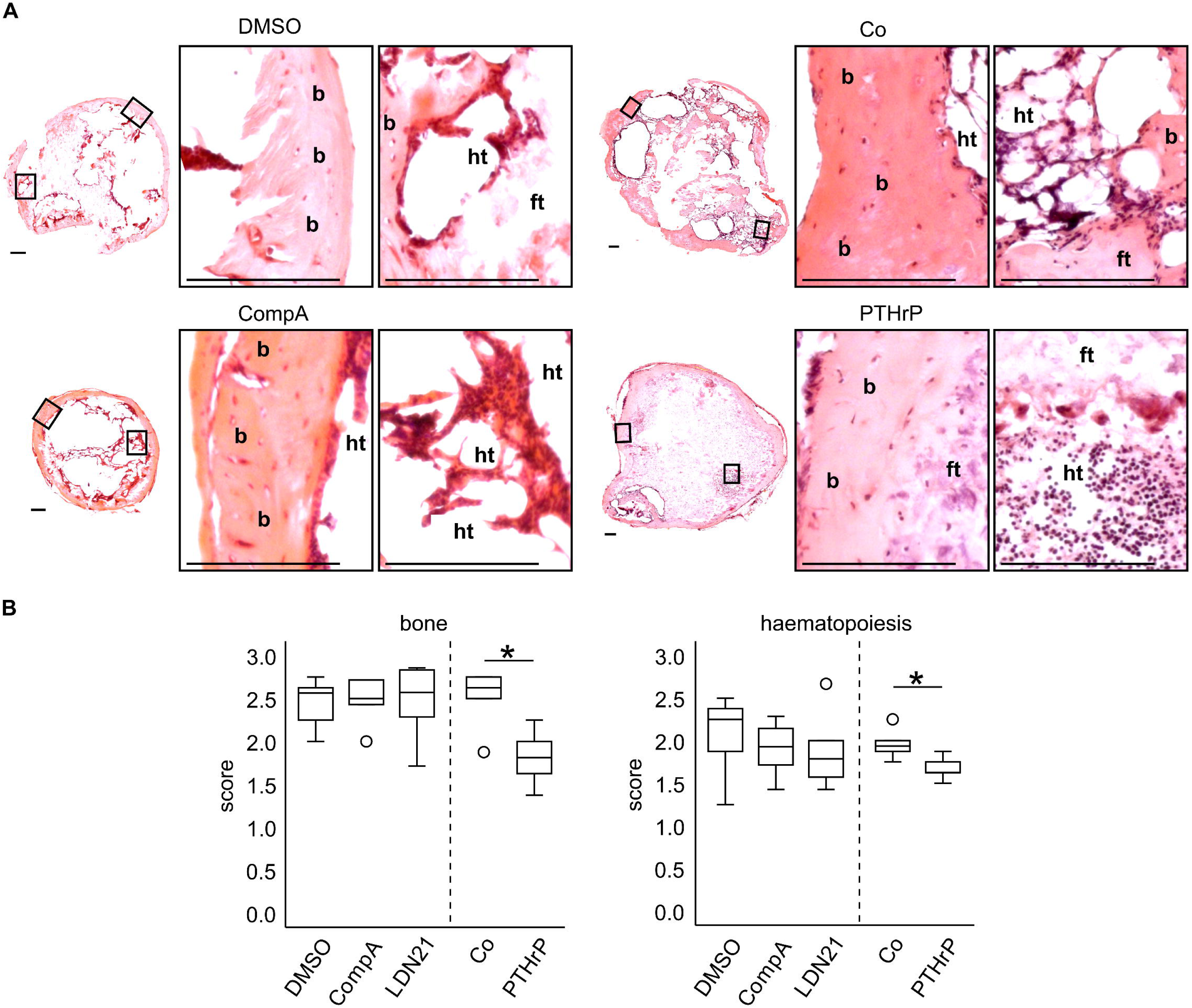
Effect of BMP pathway inhibition and PTHrP pulse treatment on bone formation and haematopoiesis of MSC-derived cartilage after 8 weeks in vivo. MSCs were subjected to chondrogenic induction in the presence of CompA, LDN21 or received PTHrP pulse treatment. After 28 days, samples were implanted in subcutaneous pouches on the back of SCIO mice for 8 weeks. (A) Paraffin sections of n=?-8 explants from n=4 MSC donors per group were stained with HE to observe bone formation and haematopoiesis. Exemplary sections are marked with b for bone, ht for haematopoiesis and ft for fibrous tissue. Scale bars represent 150 µm. (B) HE-stained sections were scored by 6 blinded scorers to define bone formation (score from 0-3) and haematopoiesis (score from 0-3). Mean scores for each treatment regimen were calculated for each scorer for statistical evaluation (n=6). Box plots were generated as described in Fig. 3. * p 0.05 Mann-Whitney U test compared to respective controls.

### ALK4/5 strongly cross-activates SMAD1/5/9 in chondrogenic MSC cultures

To address SMAD1/5/9 cross-activation in chondrogenic MSC cultures, we compared SMAD phosphorylation levels under ALK1/2/3 inhibition by LDN21 and under specific ALK4/5/7 inhibition by SB-431542 (Fig 7). Interestingly, 500 nM LDN21 reduced but did not abrogate the TGF-β-induced phosphorylation of SMAD1/9 (Fig 7A) and did not affect SMAD2/3 activation as expected (Fig 7B). This was in line with the mild overall effects of this inhibitor that we observed in chondrogenic MSC cultures. In contrast, 10 µM SB-431542 showed a strong pSMAD1/9 and pSMAD2/3 suppression. This demonstrated a strong cross-activation of SMAD1/5/9 by ALK4/5 in the presence of supraphysiologic amounts of TGF-β and only little contribution of ALK1/2/3 during MSC chondrogenesis. Thus, we propose that even a full blockage by knocking out ALK1, ALK2, ALK3, and ALK6 would not abrogate SMAD1/5/9 signalling in this system.

**Fig. 7.**
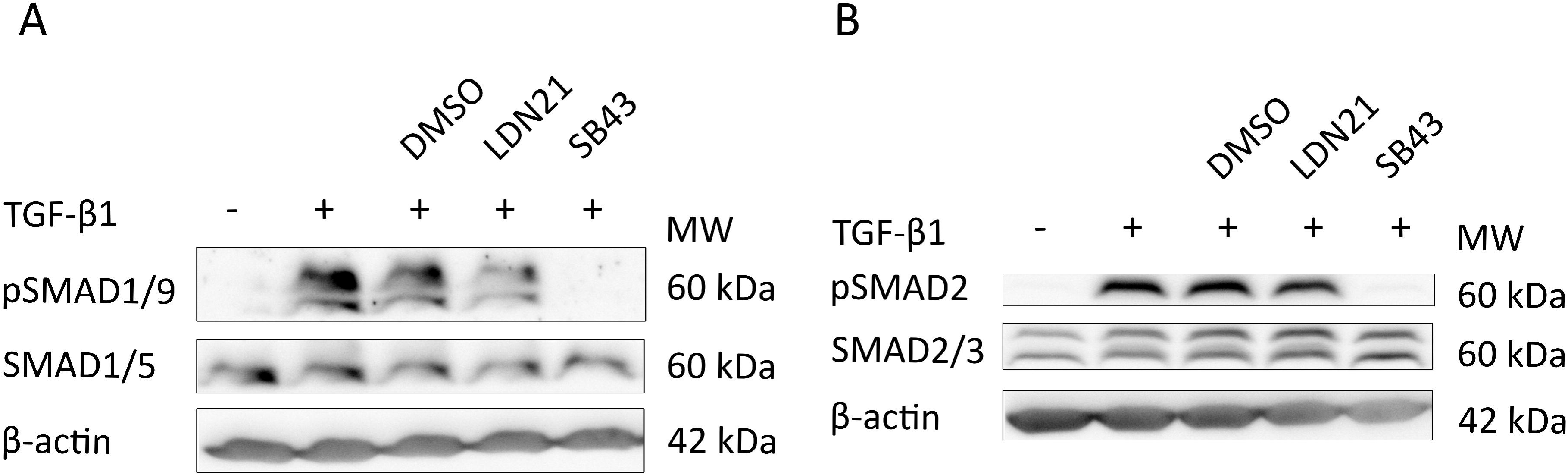
Contribution of ALK receptors to TGF--induced SMAD activation in chondrogenic MSC cultures. MSCs were subjected to chondrogenic induction in absence or presence of 10 ng/ml TGF-1, 500 nM LON 212854 (LDN21) to inhibit ALK1/2/3 or 10 µM SB-431542 (S843) to inhibit ALK4/5, or DMSO as solvent control for 30 min as indicated. (Phosphorylated) SMAD proteins were detected via Western blotting with -actin detection as internal reference. One representative blot of three experiments with three independent MSC donor populations is shown. The image was cropped for conciseness as indicated and full-length blots are presented in supplementary Fig. S7.

## Discussion

TGF-β is the only chondro-inducer that is capable of driving MSC chondrogenesis. Unfortunately, under defined standard in vitro conditions, TGF-β guides MSCs inevitably into the endochondral instead of the desired chondral lineage. Activation of the BMP-associated ALK1/2/3 receptors that signal through SMAD1/5/9 has been accused as major director misguiding this lineage choice of MSCs [12, 24]. A rather simplistic model was proposed that strictly separates this undesired TGF-β/BMP signalling pathway from the canonical TGF-β pathway that signals via ALK4/5 and SMAD2/3 and is classified as prochondrogenic and antihypertrophic [12, 26]. However, contradictive results were repeatedly reported from attempts to experimentally verify this model and evidence for extensive interlinking between the TGF-β and BMP signalling pathways is overwhelming. This stipulated us to exploit the high specificity of novel ALK1/2/3 inhibitors to decipher the prochondrogenic and prohypertrophic contributions of ALK1/2/3 activation and to assess the antihypertrophic efficacy of ALK1/2/3 inhibition by direct comparison with our current most potent antihypertrophic treatment, PTHrP pulses.

In presence of either one of the two inhibitors with the putatively highest bias to inhibit ALK1/2/3 over ALK4/5, TGF-β efficiently induced in vitro MSC chondrogenesis yielding cartilaginous tissue rich in proteoglycans and type II collagen. In sight of this data, the complete abrogation of chondrogenesis upon treatment with high-dose dorsomorphin that we observed in our previous study [11], was likely attributed to off-target inhibition of ALK4/5 and SMAD2/3 activation. Still, chondrogenic traits were also slightly underdeveloped compared to controls in the current study, which is likely to also affect the biomechanical properties, since the compressive and tensile strength of cartilage tissue is determined by the proteoglycan and collagen content. This indicated that ALK1/2/3 have some prochondrogenic activity and thereby biomechanical impact, which is, however, of little significance in the presence of strong ALK4/5 stimulation by TGF-β. Importantly, the high hypertrophy marker levels still identified the differentiation route as endochondral rather than chondral. In clear contrast to the pulsed PTHrP treatment group, slight reduction of hypertrophy markers in presence of ALK1/2/3 inhibitors was merely a bystander effect of reduced chondrogenesis. In line, the propensity of the resulting cartilaginous tissue to be remodelled into bone in vivo remained completely unchanged in the ALK1/2/3 inhibitor groups but was reduced after pulsed PTHrP treatment. This demonstrated that endochondral MSC lineage choice occurred independently of ALK1/2/3 activity which was of overall minor importance for MSC chondrogenesis.

Overall, with the current study included, we now have data on the specific application of three ALK1/2/3 inhibitors (LDN 212854, LDN 193189, compound A). In contrast to the report by Occhetta et al., the more recent data demonstrated that endochondral MSC misdifferentiation occurred independently of BMP-ALK1/2/3 activity (here and [27]). Neither ALK1/2/3 inhibitor was able to induce the desired chondral lineage shift, which is defined by strong chondrocyte formation without accompanying signs of hypertrophy, and which can be induced at least partially in vitro with pulsed PTHRP treatment or WNT inhibition or fully in vivo with heparin-coupled TGF-β [5, 6, 10]. When LDN 193189 was used in low doses that are expected to repress ALK1/2/3 but not ALK4/5, no effect on MSC chondrogenesis was observed by Franco et al. in line with the small effects observed here with compound A and LDN21 [23, 27]. Although stronger effects were reported when dorsomorphin or LDN 193189 were used in higher doses [11, 27], hypertrophy and chondrocyte markers were equally affected in line with the here presented data, even though ALK1/2/3 was expected to be 500-fold more strongly inhibited than ALK4/5. In sight of this consistent line of evidence, we conclude that BMP-ALK1/2/3 activity is no director of the MSC lineage choice during in vitro chondrogenesis, since endochondral misdifferentiation was induced by TGF-β independently of ALK1/2/3, and is likely to occur downstream of ALK4/5 and/or other pathways. This indicates, that even a full ALK1/2/3/6 knock out could not prevent endochondral misdifferentiation of TGF-β-stimulated MSCs.

LDN21 was chosen here not only as commercially available alternative to compound A, but mostly because of its over 7,000-fold higher IC50 value for ALK5 than for ALK2 in kinase assays [23]. This marks LDN21 the currently most specific ALK1/2/3 inhibitor that outperformed dorsomorphin and LDN 193189, when directly compared by kinase and cell reporter assays in vitro [23]. Also compound A was found to be more specific albeit less potent than LDN 193189 in a direct comparison, but only reached a 40-fold higher IC50 value for ALK5 over ALK2 [28]. Low inhibitor doses were necessary to prevent off-target inhibition, but the BMP activity assay in C2C12 cells confirmed a strong activity of the used LDN21 and compound A concentration. That in MSCs, by contrast, the inhibitor effects remained small, is therefore attributed to considerable ALK1/2/3-independent SMAD1/5/9 cross-activation in presence of high amounts of TGF-β, which we showed to occur downstream of ALK4/5 (Fig. 7). Taken together we conclude, that the observed low inhibitor efficacy was not a problem of underdosing, but rather caused by TGF-β-mediated cross-activation of SMAD1/5/9 via ALK4/5.

One important strength of our in vivo results compared to previous studies [12, 24] is the use of microcomputed tomography for assessing the entire original unprocessed explant, and demonstrated not only that all 46 implanted constructs including those with a cartilaginous non-mineralized core were surrounded by a calcified shell. Furthermore, micro-CT allowed the quantitative conclusion of an overall unchanged mineralization degree independently of ALK1/2/3 inhibitor pretreatment. One limit of this study is that we strongly focused our investigations on the lineage commitment during MSC chondrogenesis and can, thus, not conclude on a potential influence of ALK1/2/3 activity on the biomechanical qualities of engineered cartilage beyond those that are determined by the proteoglycan and type collagen quantities. Of note, we have previously observed that expression of several BMPs was stimulated by mechanical load in MSC-derived engineered cartilage [29], and the importance of ALK1/2/3 induction for the mechanoresponse and mechanoadaptation of tissue properties remains an interesting question.

We here showed for the first time that ALK4/5 strongly cross-activates SMAD1/5/9 during MSC chondrogenesis where supraphysiologic TGF-β concentrations are routinely applied. Together with the negligible role of ALK1/2/3 activation for MSC chondrogenesis, this invites to adapt the concept of dual TGF-β activity and test the still simplistic hypothesis of a prochondrogenic ALK4/5-SMAD2/3 activity contrasted by a prohypertrophic ALK4/5-SMAD1/5/9 activity. In support of this hypothesis, we previously observed that sulfation of glycosaminoglycan hydrogels which can instruct cell fate and chondral versus endochondral lineage decision of MSCs in vivo, also appeared to differentially affect SMAD1/5/9 and SMAD2/3 signalling according to reporter assays [5]. Thus, the chondral versus endochondral lineage switch appeared to coincide with a shift in SMAD preference. However, also WNT and hedgehog signalling were affected, indicating an intimate interaction of all prohypertrophic signalling pathways that regulate MSC chondrogenesis. Beyond the importance of SMAD1/5/9 for the hypertrophic development during MSC chondrogenesis, also the occurrence and significance of mixed R-SMAD complexes needs to be addressed. Unfortunately, such experiments are complicated by the current unavailability of specific SMAD inhibitors and the incompatibility of MSC in vitro chondrogenesis with genetic engineering techniques that could facilitate long-term modification of SMAD levels.

## Conclusion

We here demonstrated that endochondral MSC misdifferentiation during in vitro chondrogenesis occurs independently of ALK1/2/3 signalling, and ALK1/2/3 activation was shown dispensable for prochondrogenic signalling. Unlike suggested before [12, 24], induction of ALK1/2/3 was no decisive misfunction of the chondro-inducer TGF-β leading to hypertrophy, and ALK1/2/3 inhibition was not capable to program MSCs into stable chondrocytes that self-sustain their phenotype in vivo. We therefore propose to revise the current concept of a prohypertrophic BMP-ALK1/2/3, WNT, hedgehog signalling network during TGF-β-driven in vitro chondrogenesis of MSCs, into a TGF-β-ALK4/5, WNT, hedgehog signalling network which interactively regulates endochondral misdifferentiation of MSCs. Further deciphering of the prohypertrophic TGF-β activity is still needed including the assessment of the exact role of SMAD1/5/9 activation, mixed R-SMAD complexes and of further non-canonical TGF-β pathways that may contribute to hedgehog and WNT activity in vitro but not when TGF-β is coupled to heparin in an in vivo setting. This will help to overcome the high resilience of the prohypertrophic signalling network in MSCs and to translate in vitro engineered MSC-based neocartilage replacement tissue to clinical cartilage regenerative therapy.

## Declarations

### Ethical approval

Bone marrow samples were obtained after approval by the Ethics Committee on Human Experimentation of the Medical Faculty of Heidelberg University (S-117/2014 ‘Molekulare Charakterisierung hypertropher Differenzierungswege und ihre Vermeidung bei der in vitro Chondrogenese humaner mesenchymaler Stammzellen’ approved in May 2014, S-845/2019 ‘Gezielte Steuerung hypertropher Differenzierungswege zur Verbesserung der in vitro Chondrogenese humaner mesenchymaler Stammzellen’ approved in December 2019) and in agreement with the Helsinki Declaration of 1975 in its latest version.

All animal experiments were approved by the local animal experimentation committee (G-210/18) and carried out in accordance with European Laboratory Animal Science guidelines.

### Consent for publication

n/a

### Author Contributions

SD: Experimental design, data interpretation, manuscript writing, final approval of manuscript. SID: Conception and design, collection of data, data analysis and interpretation, manuscript writing, final approval of manuscript. SN: Provision of study material, collection of data, data analysis and interpretation, final approval of manuscript. SvS: Collection of data, manuscript preparation, final approval of manuscript. CM: Provision of study material, final approval of manuscript. WR: Conception and design, financial support, administrative support, data interpretation, manuscript writing, final approval of manuscript.

## Acknowledgements

The authors thank Nina Hofmann, Birgit Frey, and Felina Zahnow for their excellent technical help.

## Data Availability Statement

The raw data supporting the conclusions of this manuscript will be made available by the authors, without undue reservation, to any qualified researcher upon request.

