## Supplementary figures and images for "Mesenchymal stromal cell chondrogenesis under ALK1/2/3-specific BMP inhibition: A revision of the prohypertrophic signalling network concept"

### Suppl. Figure 1

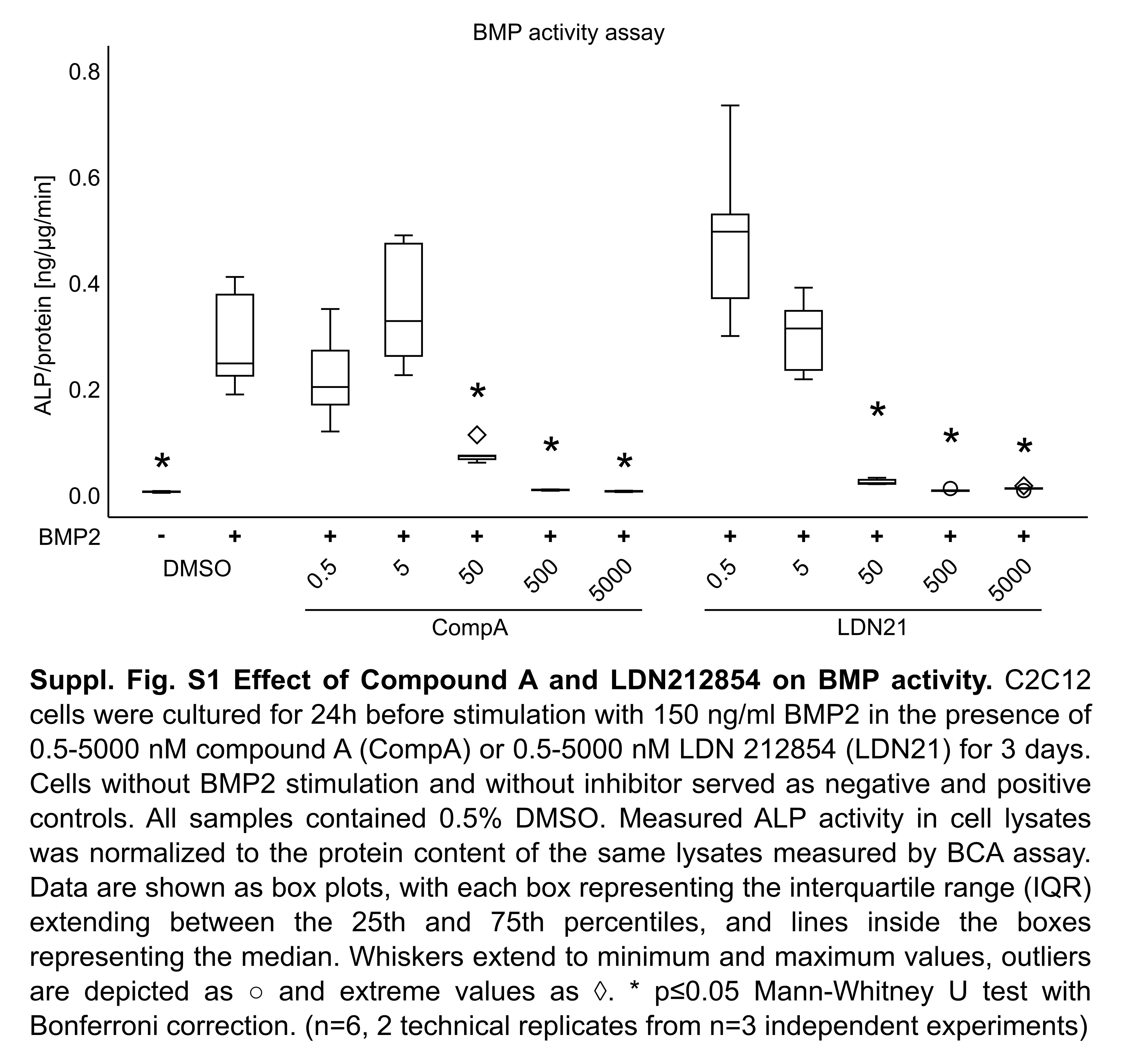

### Suppl. Figure 2

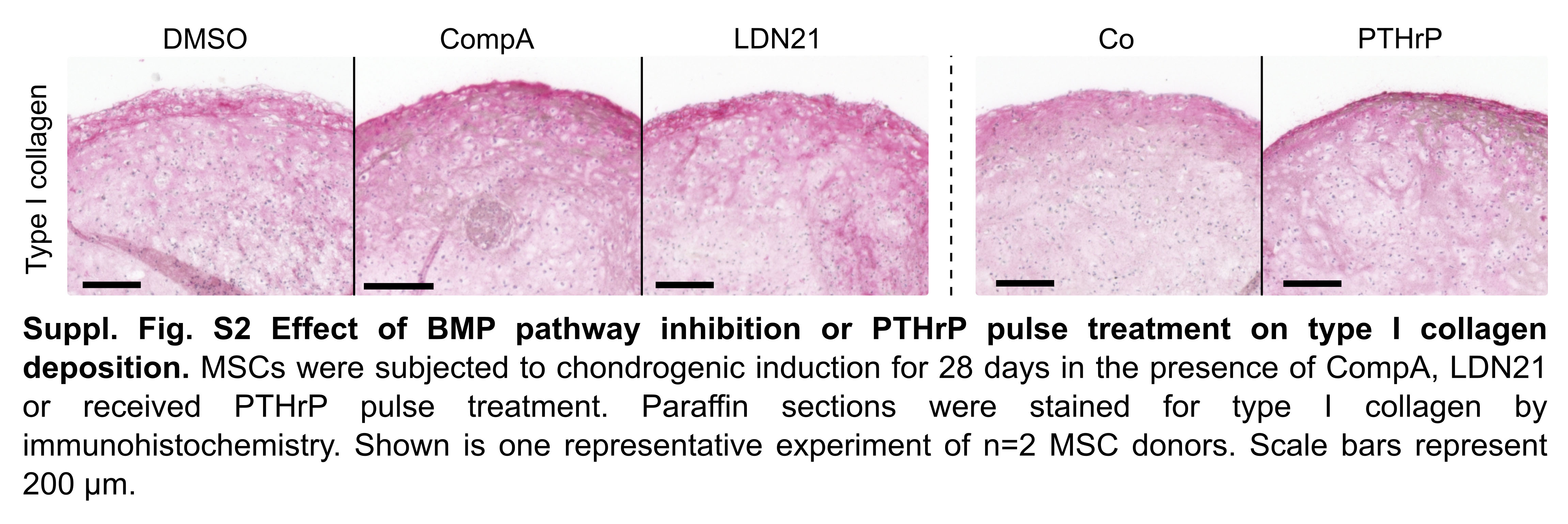

### Suppl. Figure 3

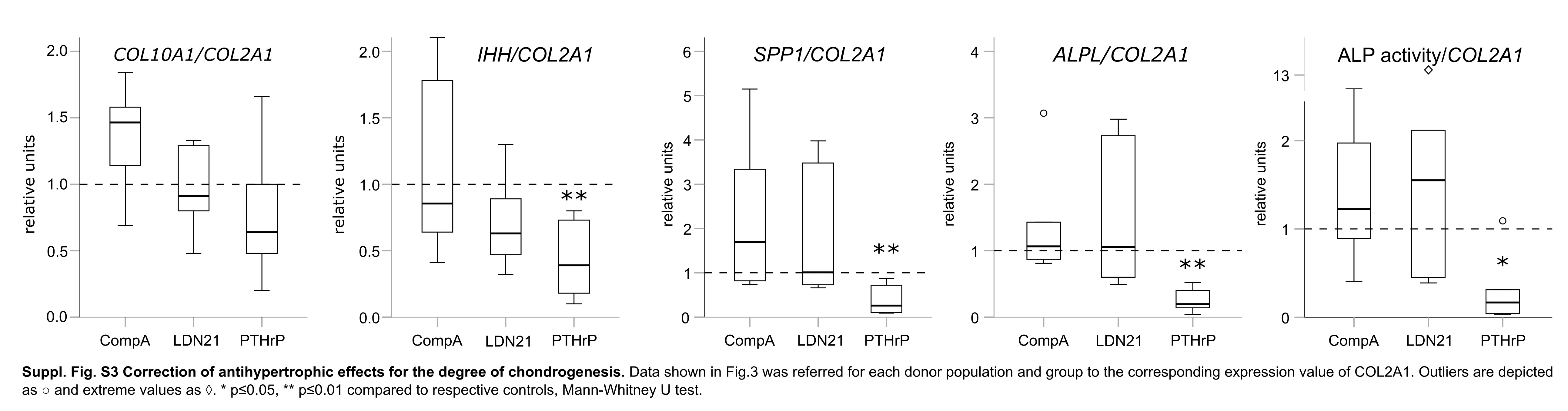

### Suppl. Figure 4

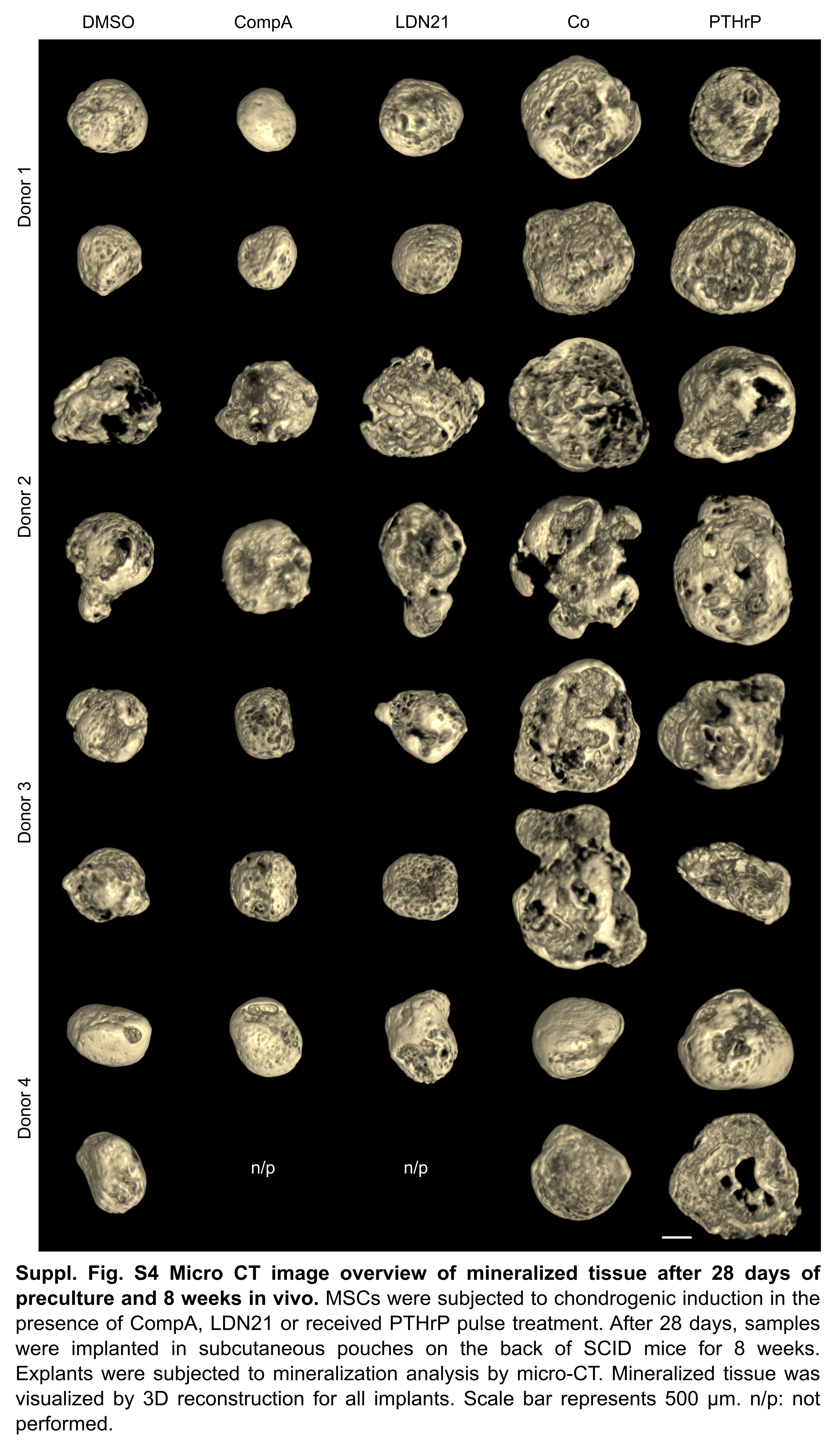

### Suppl. Figure 5

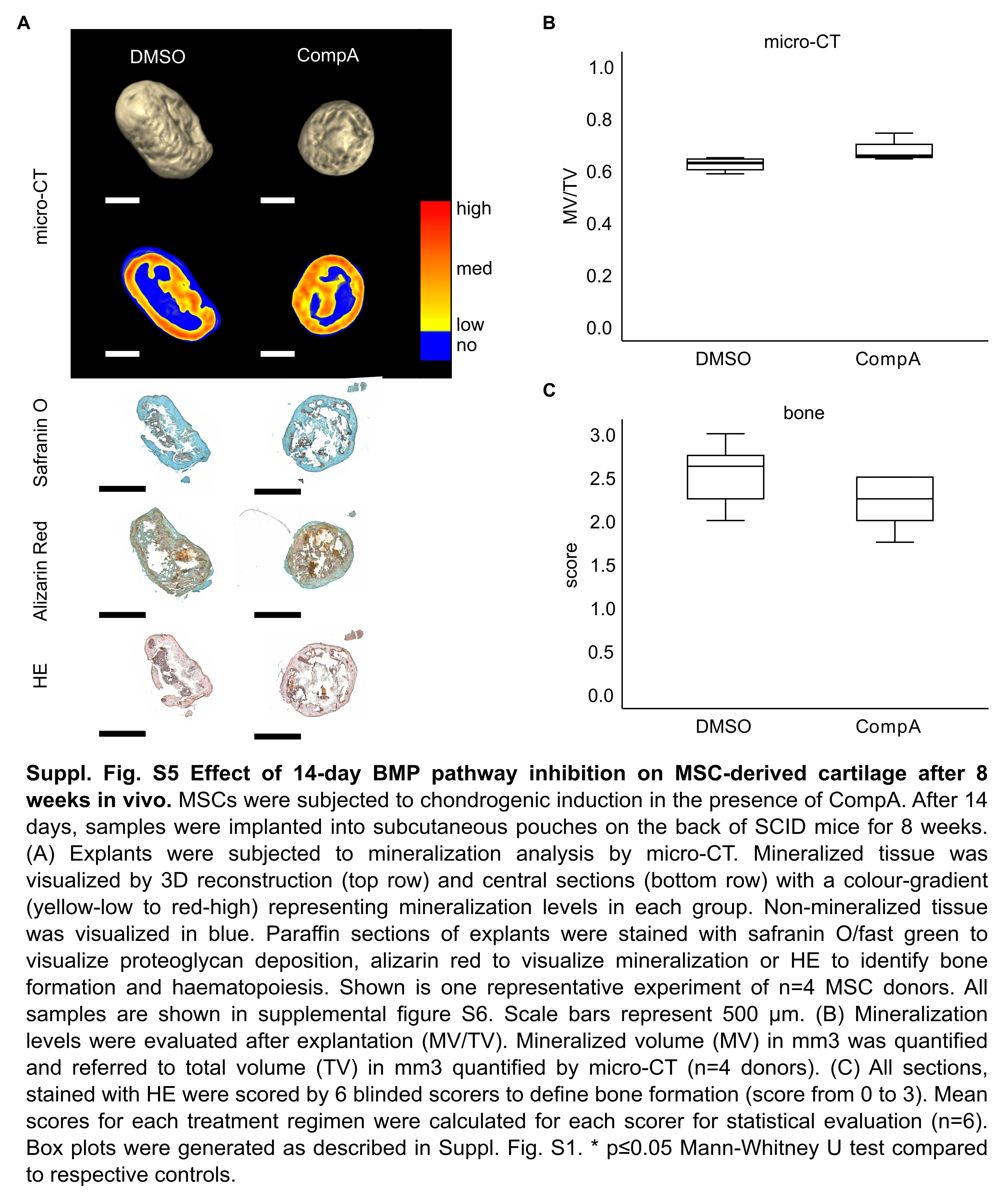

### Suppl. Figure 6

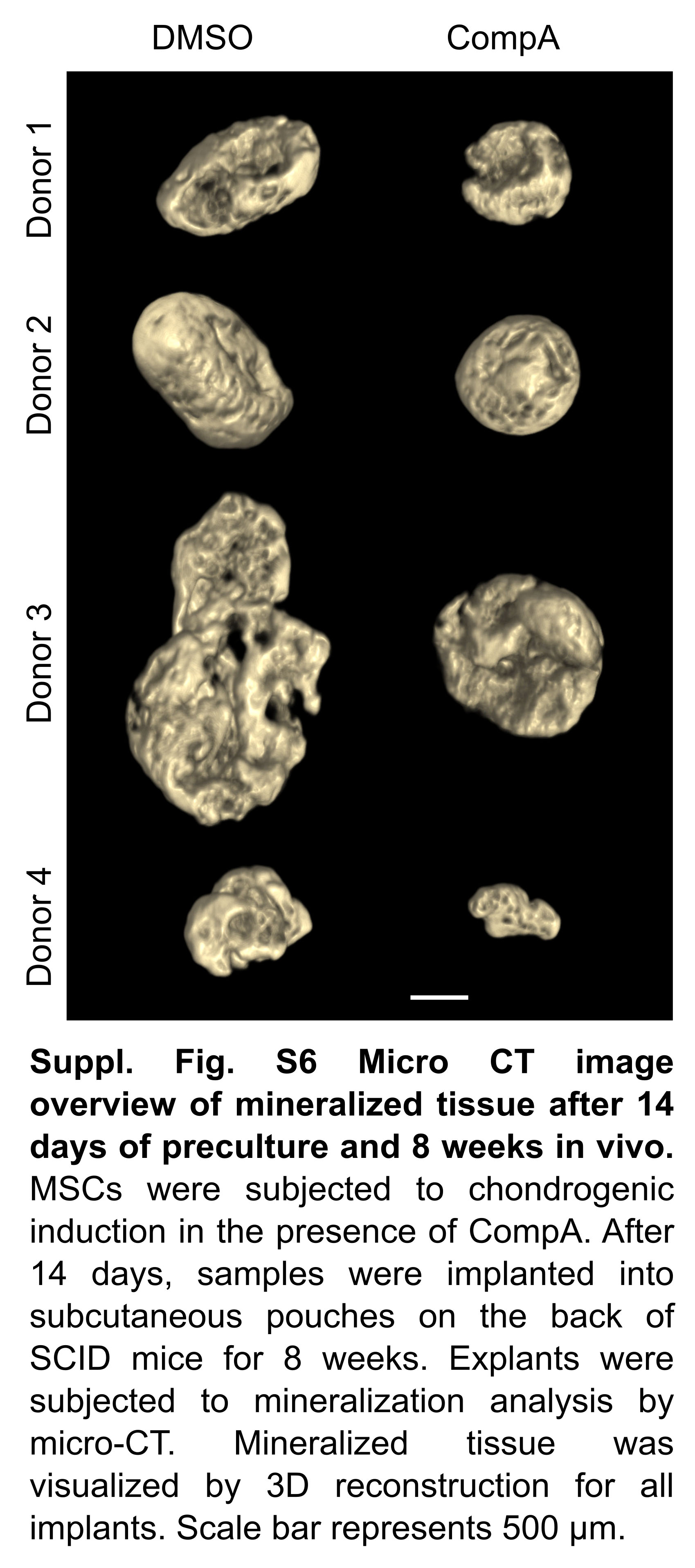

### Suppl. Figure 7

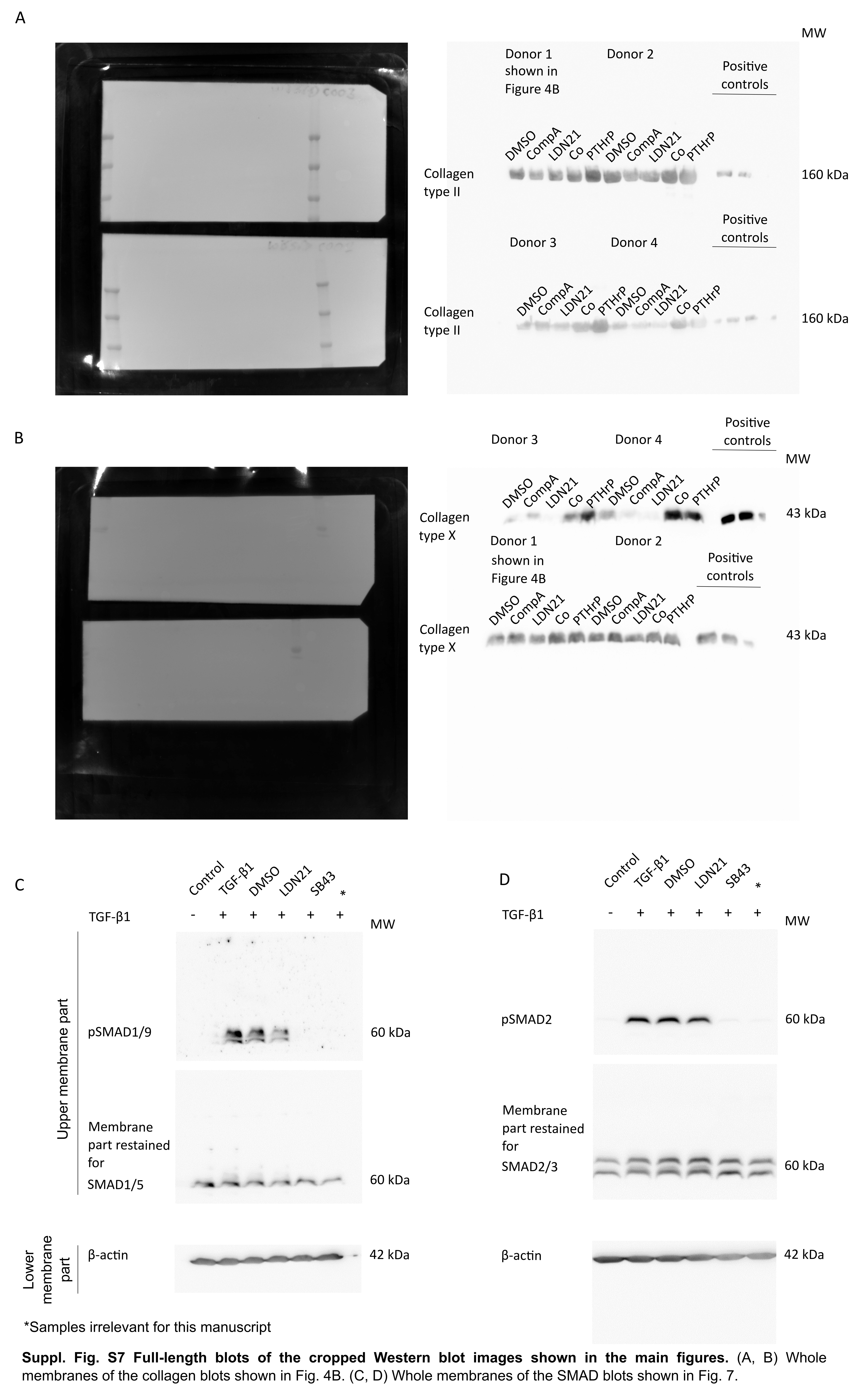
